# SP1 affects the migration and invasion of laryngeal cancer cells by regulating methylation of claudin 4 promoter region through the p-JNK pathway

**DOI:** 10.1101/2022.07.20.500879

**Authors:** Jinsong Ni, Hailian Shi, Kaiyue Peng, Ya Liu, Yafang Liu

## Abstract

Claudin 4 (CLDN4) is abnormally expressed in various tumors, but the mechanisms controlling CLDN4 expression in laryngeal cancer are poorly understood. Here, we describe hypermethylation of the promoter region of the CLDN4 gene and a positive correlation between the expression of CLDN4 and transcription factor Sp1 in laryngeal carcinoma. Specifically, CLDN4 expression was downregulated when laryngeal cancer cells were treated with Sp1 transcription factor inhibitor MTM. Immunohistochemical staining showed that while the expression of CLDN4 and p-JNK was negatively correlated, p-JNK positively correlated with MMP-2 and MMP-9 expression. The expressions of p-JNK, MMP-2, and MMP-9 were significantly downregulated in HEp-2 cells transfected with CLDN4, decreasing migration and invasiveness of HEp-2 cells. Thus, we show that SP1-mediated hypermethylation of the CLDN4 promoter region may activate the JNK signaling pathway and increase MMP-2/MMP-9 expression, which can then affect the migration and invasiveness of laryngeal cancer cells. These results contribute to our understanding of the regulatory mechanisms of CLDN4 in laryngeal cancer and identify the Sp1-CLDN4-jnk signaling pathway-MMP-2/MMP-9 axis as a potential target for the treatment of laryngeal cancer.

## Introduction

Claudins are a family of transmembrane proteins in tight junctions divided into two groups, namely, classical claudins (1 – 10, 14, 15, 17, 19) and non-classical claudins (11 – 13, 16, 18, 20 – 24), based on sequence similarity [1]. As a member of the claudin family, CLDN4 is abnormally expressed in a variety of human tumor tissues [2,3]. We previously showed that demethylation of claudin 4 (CLDN4) DNA could inhibit the migration and invasion of laryngeal squamous cell carcinoma [4]. DNA methylation can affect the transcriptional inactivation of tumor suppressor genes in two ways. First, methylated DNA hinders transcription activators’ binding and recognition sites, e.g., binding sites for transcription factors such as NF-KB, SP1/SP3, and Sp1 contain CpG islands. Their methylation can inhibit the binding of transcription factors [5,6,7]. Second, methylated DNA can recruit methyl-CpG-binding proteins (Mbps), which inhibit gene transcription, and the methyl-CpG binding domain (MBD) may be involved in this process. MBD has five family members, namely, MeCP2, MBD1, MBD2, MBD3, and MBD4. While all have MBD domains, only MeCP2, MBD1, and MBD2 have a transcriptional-repression domain, which inhibits gene transcription by binding to specific transcription inhibitors [8,9]. Previously, we have reported that the expression of cldn4 negatively correlated with the expression of MeCP2 [4].

Abnormalities in the mitogen-activated protein kinase (MAPK)-JNK pathway are usually involved in the occurrence and progression of cancer, and many studies have confirmed that the MAPK-JNK signaling pathway is related to cancer metastasis and invasion [10]. Hence, we analyzed whether the transcription factor Sp1 plays a role in DNA methylation in laryngeal squamous cell carcinoma and if it is associated with changes in CLDN4 expression. We also investigated the possible effects of CLDN4 on the MAPK-JNK signaling pathway in laryngeal squamous cell carcinoma.

## Results

### 1.1 Expression of and correlation between CLDN4 and Sp1 in laryngeal carcinoma

The methylation status of the SP1 site in the promoter region of mir-23a-27a-24-2 can promote proliferation and inhibit early apoptosis in laryngeal cancer cells [11]. To determine the link between SP1 and CLDN4, we performed immunohistochemical staining for Sp1 and clnd4 in 84 laryngeal squamous cell carcinoma tissue samples and 76 adjacent tissue samples. Among laryngeal carcinoma samples, CLDN4 expression was positive in 54 cases and negative in 30 cases, while SP1 expression was positive in 79 cases and negative in 5 cases. Next, CLDN4 was expressed in the cell membrane and cytoplasm of laryngeal cancer tissue, and its intensity in laryngeal carcinoma was higher than that in adjacent tissues. In contrast, Sp1 was present in the nucleus and showed stronger expression in cancer tissues (Figure 1 A-D). Among the 76 adjacent tissue samples tested, 70 were positive for CLDN4 while 6 were negative; SP1 expression was positive in 60 cases and negative in 16 cases (Table 1). Importantly, 53cases were positive for both CLDN4 and Sp1, 1 case was positive for CLDN4 and negative for Sp1, 26 cases were negative for CLDN4 but positive for Sp1, and 4 cases were negative for CLDN4 and Sp1. Spearman’s correlation analysis showed that the expression of CLDN4 was correlated with the expression of Sp1 (Table 2).

**Table 1.** Expression of CLDN4 and Sp1 in laryngeal carcinoma and adjacent tissues

|  | **Total** | **Positive** | **Negative** | **Positive rate** |
| --- | --- | --- | --- | --- |
| **CLDN4 in laryngeal carcinoma** | 84 | 54 | 30 | 64.3% |
| **CLDN4 in paracancerous tissues** | 76 | 70 | 6 | 92.1% |
| **SP1 in laryngeal carcinoma** | 84 | 79 | 5 | 94.0% |
| **SP1 in**<br>**paracancerous**<br>**tissues** | 76 | 60 | 16 | 78.9% |

**Table 2.** Correlation between CLDN4 and Sp1 expression

| CLDN4 | MMP2 |  | spearman | P 值 |
| --- | --- | --- | --- | --- |
| | Positive | Negative | $r_s$ | |
| Positive | 53 | 1 | 0.233 | <0.05 |
| Negative | 26 | 4 |  |  |

**Figure 1.**
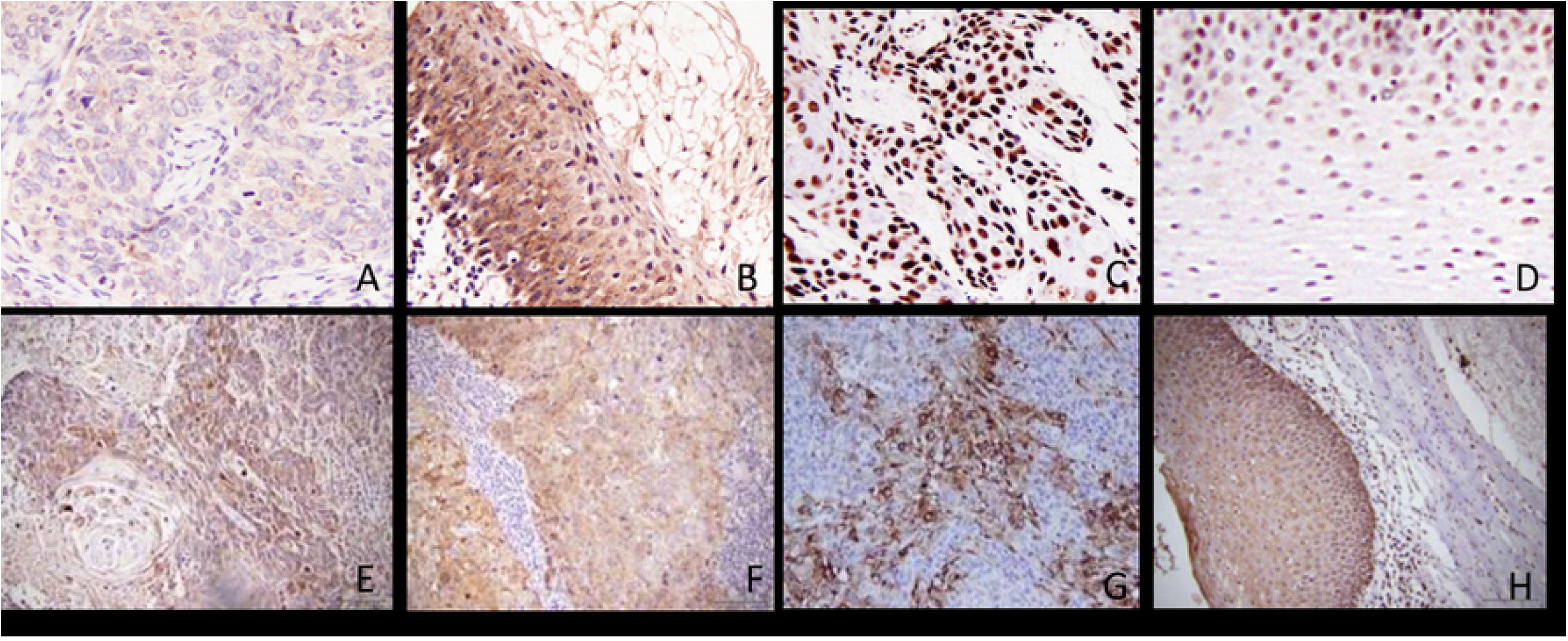
(A) Immunohistochemical staining of CLDN4 in laryngeal squamous cell carcinoma was weakly expressed in cell membrane and cytoplasm (B) Immunohistochemical staining of CLDN4 in adjacent tissues showed moderate expression in cell membrane and cytoplasm (C) The immunohistochemical staining of SP1 was strongly expressed in the nucleus of laryngeal squamous cell carcinoma (D) In the tissues adjacent to laryngeal carcinoma, SP1 was moderately expressed in the nucleus (E) P-JNK was weakly positive in laryngeal carcinoma by immunohistochemical staining (F) MMP2 was weakly positive in laryngeal carcinoma by immunohistochemical staining (G) MMP9 was partially positive in laryngeal carcinoma by immunohistochemical staining (H) P-JNK was moderately positive in tissues adjacent to laryngeal carcinoma by immunohistochemical staining

### 1.2 Detection of methylation sites in the promoter region of CLDN4 in laryngeal carcinomaa

The protein encoded by SP1 is a zinc finger transcription factor that binds to GC-rich motifs on many promoters. We used CpG island search software to analyze the sequence of the promoter region of the *cldn4* gene. There were many CpG islands in the *cldn4* gene sequence, including at the binding site of the transcription factor Sp1 (from transcription start site + 22bp to 356bp). Hence, primers were designed for the 335 bp gene fragment that contained 21 CpG sites, such that the framed region represents primer positions, the middle part is the sequence to be tested, and the red parts are the CpG site (Figure 2A). To further analyze the methylation status of the CLDN4 promoter in laryngeal cancer cells, we cultured laryngeal squamous cell carcinoma (Hep-2 cell line) and treated it with the demethylation 5-aza-dC. We detected methylation status using bisulfite sequencing (BSP). Sequencing revealed that in the control group and compared to the original *cldn4* gene sequence, the amplified sequence contained 21 CpG sites with an average 21 possible sites of methylation, implying an average methylation probability of 100%. Importantly, these results indicate hypermethylation of the CLDN4 promoter region in HEp-2 cells. Compared to the original cldn4 gene sequence in the Administration group, the amplified sequence contained 21 CpG sites, with an average of 15 methylated sites and an average methylation probability of 71.4%. Further, compared to the control group, 19 sites were demethylated to varying degrees, with an average demethylation probability of 28.6% (Figure 2 B-D). These results imply that 5aza-dC treatment demethylated CLDN4 hypermethylation in HEp-2 (laryngeal cancer) cells.

**Figure 2.**
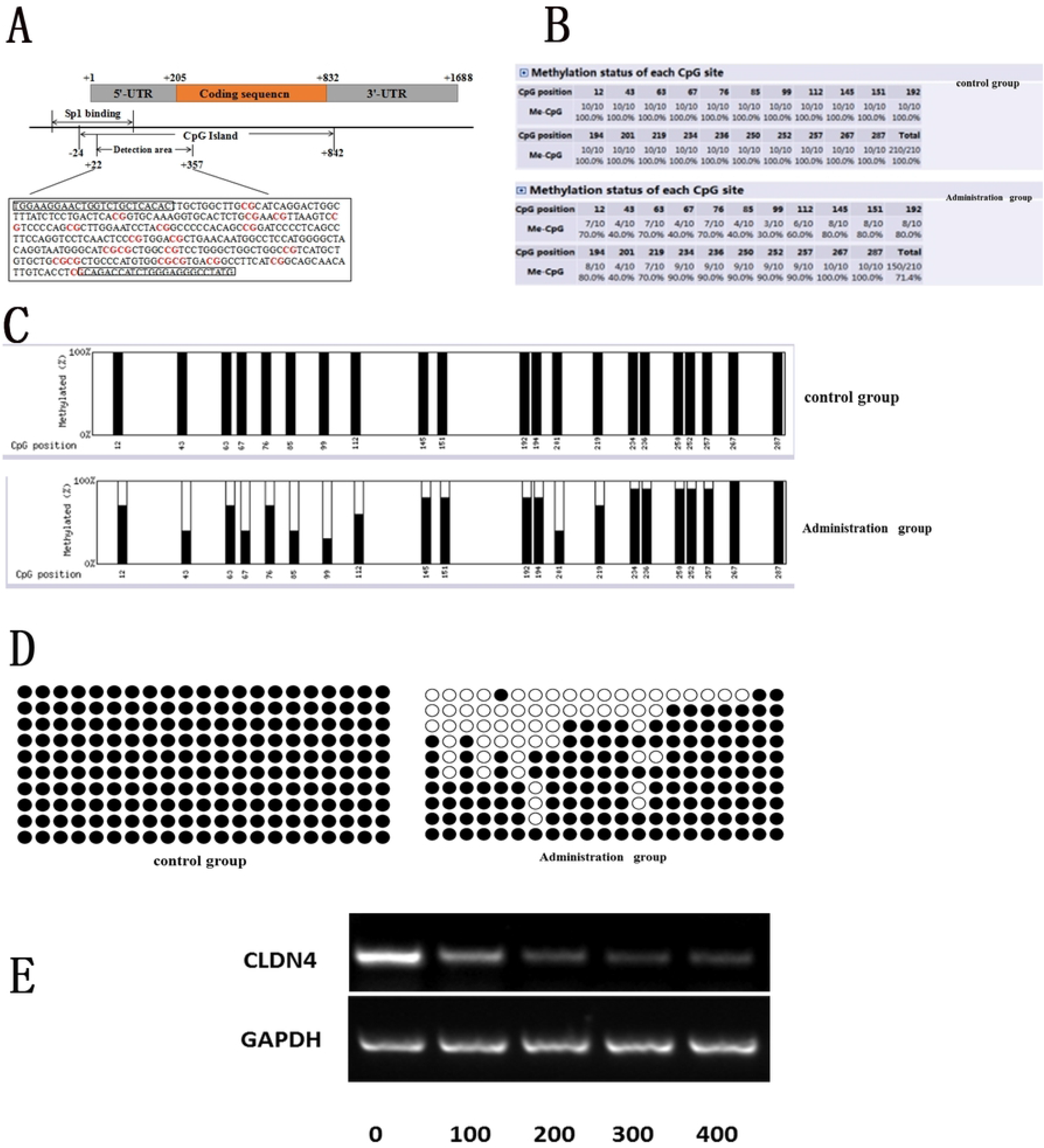
(A) CpG Island Search software analyzed the promoter region sequence of CLDN4 gene and showed 21 red CpG loci in the promoter region sequence of CLDN4 gene (B) Methylation probability of each methylation site (C) Bar graph of methylation probability of each methylation site (D) Bead diagram of methylation probability of each methylation site (E)HEp-2 cells were treated with 0, 100, 200, 300, or 400 nmol/L of the SP1 inhibitor, MTM, for 36 hours, and CLDN4 expression was detected by RT-PCR.

### 1.3 Effect of SP1 on CLDN4 expression in HEp-2 cells

HEp-2 cells were treated with 0, 100, 200, 300, or 400 nmol/L of the SP1 inhibitor, MTM, for 36 hours, and CLDN4 expression was detected by RT-PCR. All tested concentrations of MTM could downregulate CLDN4 expression in HEp-2 cells, and the maximum inhibitory effect was seen at 400 nmol/L, as shown in Figure 2E.

### 2.1 Expression and correlation of CLDN4, p-JNK, MMP-2, and MMP-9 in human laryngeal carcinoma

Changes in CLDN4 expression in cancer cells can affect their proliferation by regulating the p-JNK pathway; however, the specific mechanism is not established [12]. One study demonstrated that inhibition of the JNK signaling pathway might be related to decreased enzyme activity and protein level of matrix metalloproteinases (MMP) - 2 and MMP-9, which then inhibits cell metastasis [13]. In contrast, another study suggested that inhibiting the expression of MMP-2 and MMP-9 can activate the JNK pathway, thereby affecting the migration and invasiveness of cancer cells [14]. Therefore, we analyzed the expression of CLDN4, p-JNK, MMP-2, and MMP-9 in 80 samples of human laryngeal carcinoma and 65 adjacent tissue samples using immunohistochemical staining and evaluated the correlation between the expression of CLDN4 and the expression of p-JNK, MMP-2, and MMP-9 using Spearman correlation analysis (Figure 1 E-H). Correlation analysis revealed that CLDN4 expression negatively correlated with that of p-JNK, MMP-2, and MMP-9 (P < 0.05) and that p-JNK expression positively correlated with MMP-2 and MMP-9 expression (P < 0.05; Table 3.1-3.5).

**Table 3.1.**
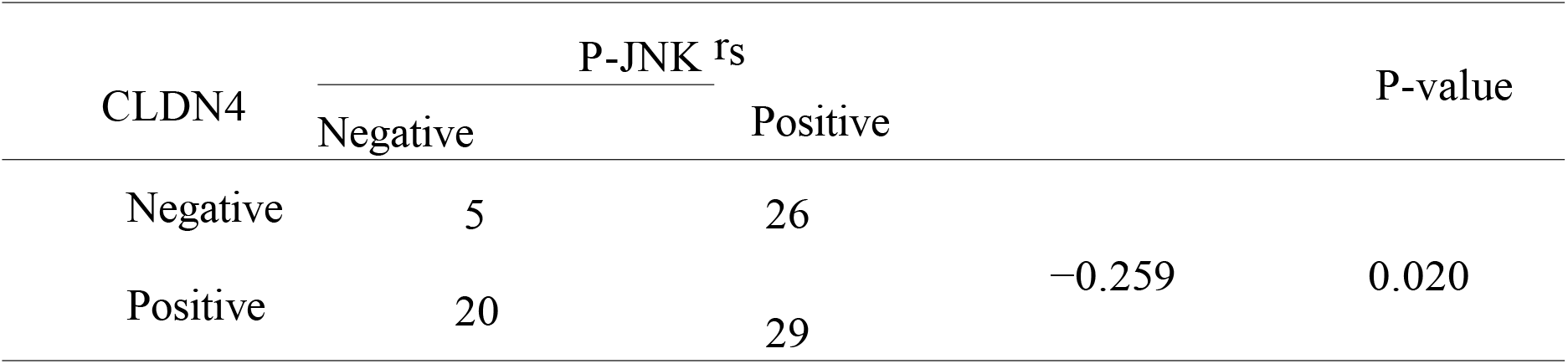
Correlation between CLDN4 and p-JNK expression

**Table 3.2.**
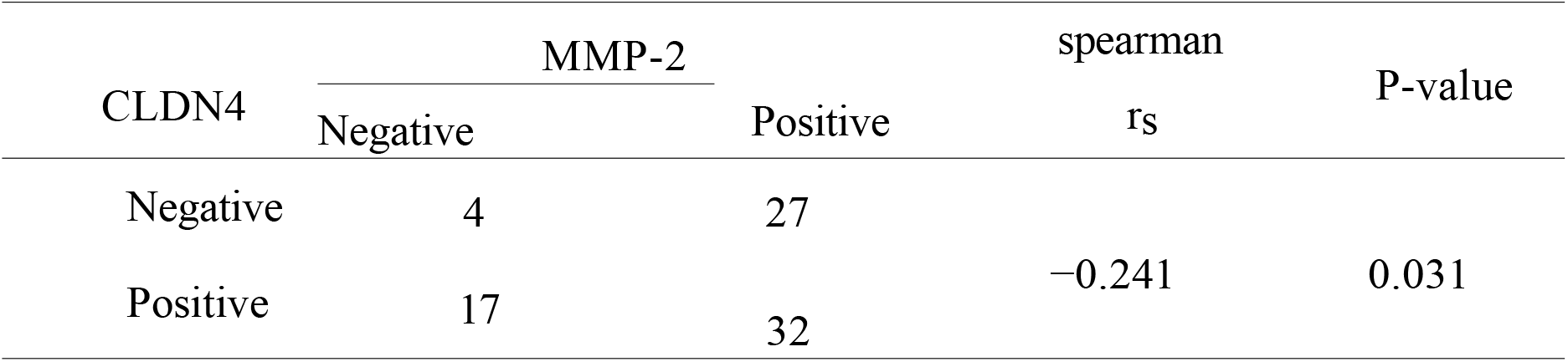
Correlation between CLDN4 and MMP-2 expression

**Table 3.3.**
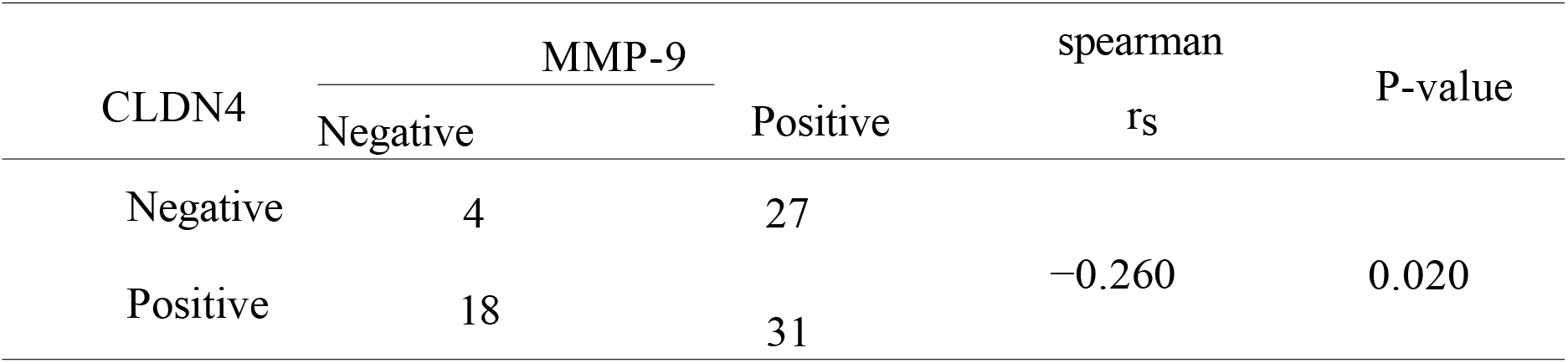
Correlation between CLDN4 and MMP-9 expression

**Table 3.4.**
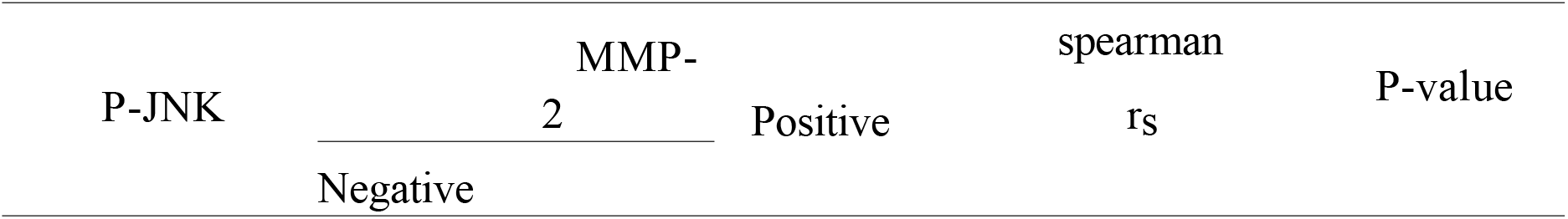
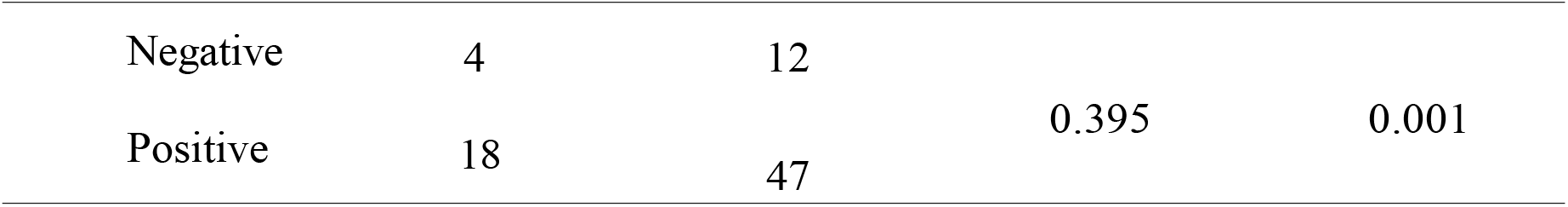
Correlation between P-JNK and MMP-2 expression

| P-JNK | MMP-2 |  | spearman | P-value |
| --- | --- | --- | --- | --- |
| | Negative | Positive | $r_s$ | |
| Negative | 4 | 12 | 0.395 | 0.001 |
| Positive | 18 | 47 |  |  |

**Table 3.5.** Correlation between P-JNK and MMP-9 expression

| P-JNK | MMP-9 |  | spearman | P-value |
| --- | --- | --- | --- | --- |
| | Negative | Positive | $r_s$ | |
| Negative | 15 | 10 | 0.354 | 0.001 |
| Positive | 7 | 48 |  |  |

### 2.2 Effects of CLDN4 overexpression on migration and invasiveness of HEp-2 Cells

We used RT-PCR and Western blotting to characterize the expression of CLDN4 in human laryngeal cancer HEp-2 cells and show that the expression of CLDN4 mRNA and protein was lower in Hep-2 cells compared to HeLa cells in the control group (P < 0.05; Figure 3 A, B). Next, we induced stable overexpression of CLDN4 in HEp-2 cells by transfection and verified significant upregulation in expression (P < 0.05) compared to the empty group, using RT-PCR and Western blotting (Figures 3 C and D). Subsequently, we evaluated the effect of CLDN4 overexpression on the migration ability of HEp-2 cells using the scratch assay and found that, compared to the empty group, the migration ability of cells overexpressing CLDN4 was significantly lower (P < 0.05; Figure 3E and F). We also tested the effect of CLDN4 overexpression on the invasiveness of HEp-2 cells using a Transwell set-up. Compared to the empty group, the invasiveness of cells overexpressing CLDN4 was significantly lower (P < 0.05; Figure 3 G and H).

**Figure 3.**
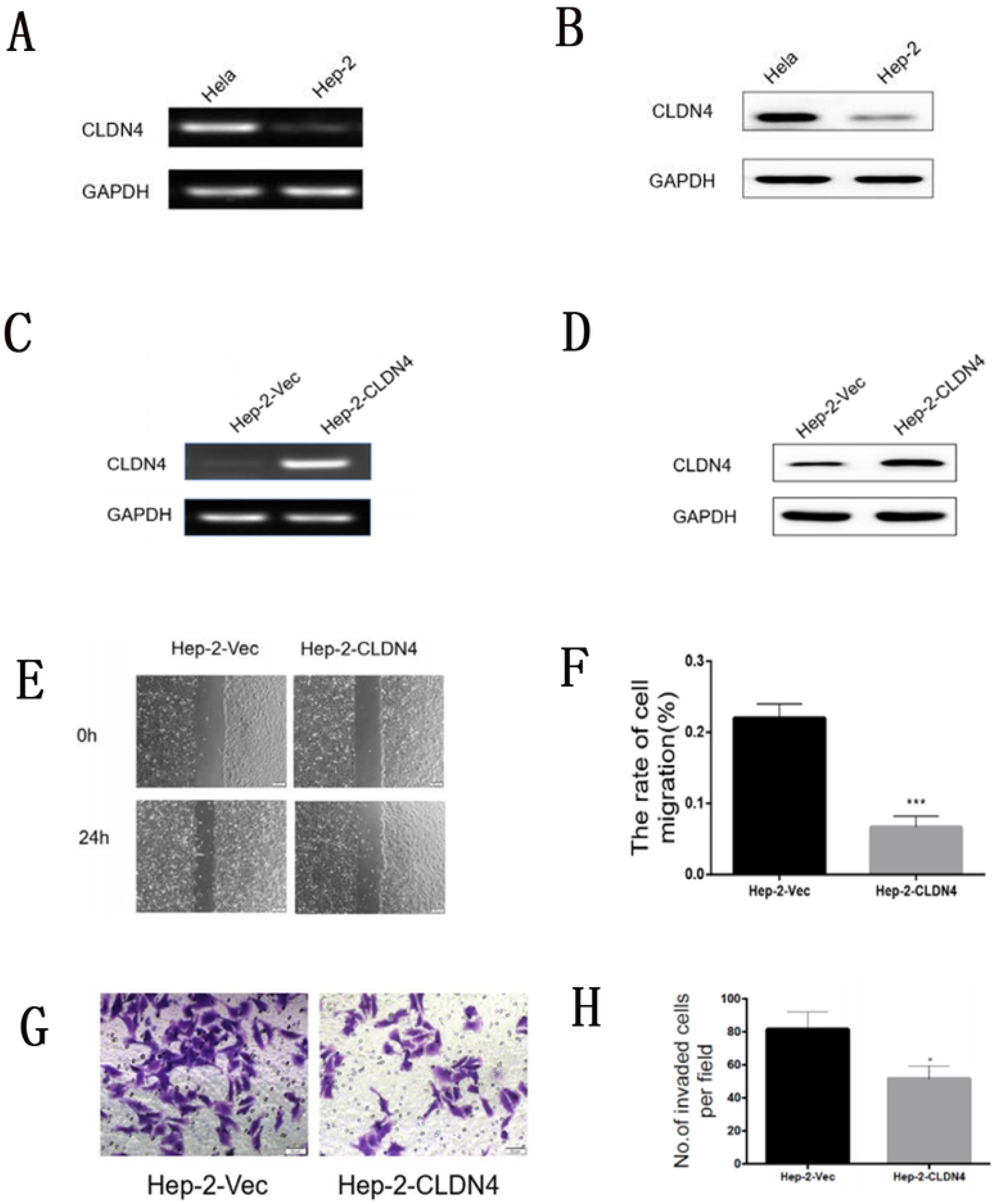
(A) RT-PCR showed that the mRNA level of CLDN4 in HEp-2 cells was lower than that in HeLa cells (B) Western blot showed that the protein level of CLDN4 in HEp-2 cells was lower than that in HeLa cells (C) RT-PCR showed that the mRNA level of CLDN4 was up-regulated in the transfection group (D) Western-blot showed that the protein level of CLDN4 was up-regulated in the transfection group (E-F) Changes of migration ability of HEp-2 cells transfected with CLDN4 in vitro (G-H) Changes of invasion ability of HEp-2 cells transfected with CLDN4 in vitro.

### 2.3 Effect of CLDN4 on the expression of p-JNK, MMP-2, and MMP-9 in HEp-2 cells

Both migration and invasiveness of HEp-2 cells decreased significantly upon CLDN4 overexpression. We speculated that it might be related to the JNK-MAPK signaling pathway. Therefore, we analyzed the expression of ERK, p-ERK, p38, p-p38, JNK, and p-JNK proteins in the Empty and CLDN4 overexpression groups using Western blotting, which demonstrated that compared to the Empty group, the expression of p-JNK in the overexpression CLDN4 group was significantly lower (P < 0.05). The expression of JNK, ERK, p-ERK, p38, and p-p38 did not change significantly (Figures 4 A and B). Similarly, the expression of MMP-2 and MMP-9 were detected by Western blot, and compared to the Empty group, MMP-2 and MMP-9 expression in CLDN4 overexpression group was significantly downregulated (P < 0.05; Figure 4 C and D).

**Figure 4.**
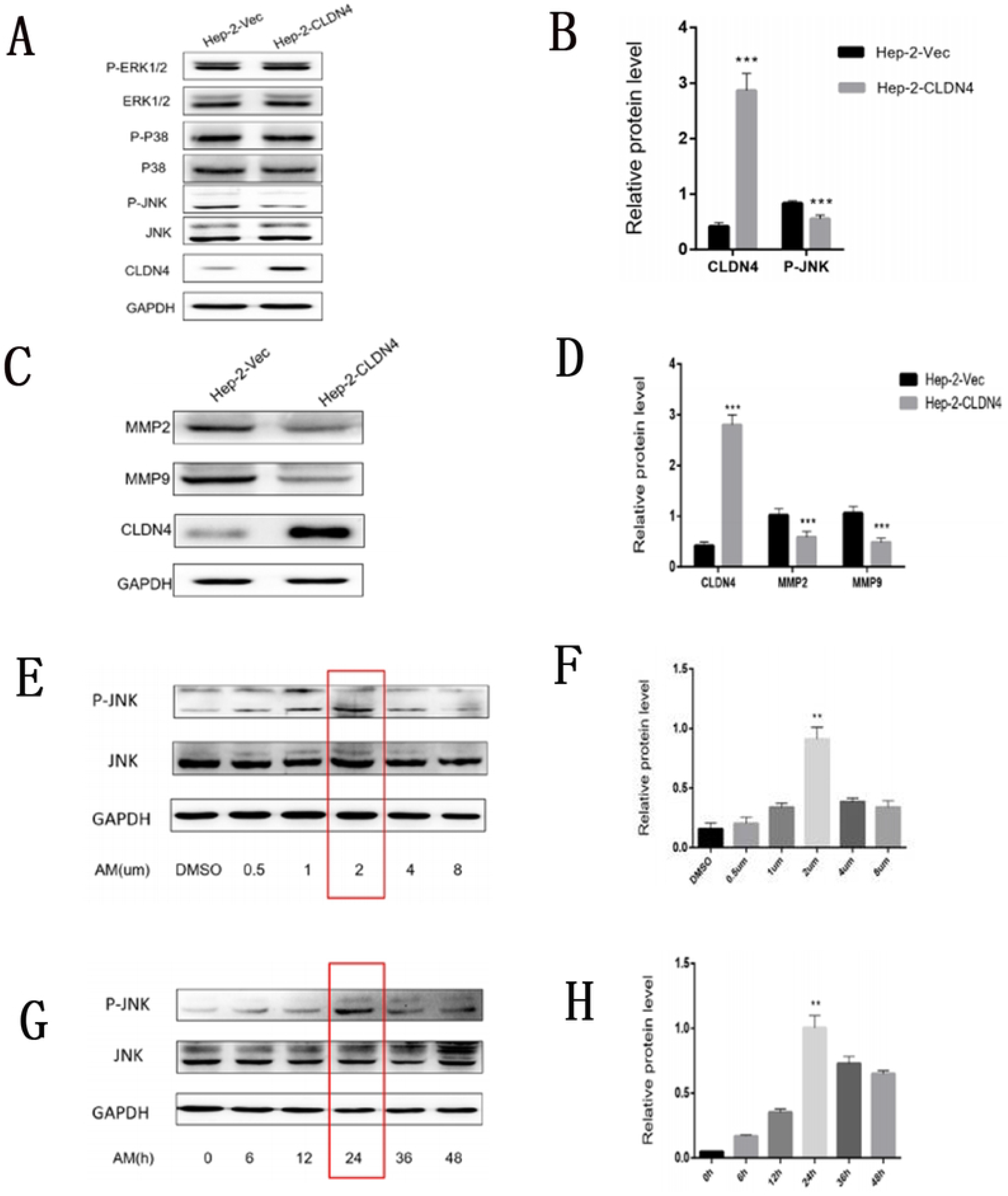
(A) Expression of MAPK-JNK signaling pathway related proteins in HEp-2 cells transfected with CLDN4 (B) The histogram showed that the expression of p-JNK was down-regulated in HEp-2 cells transfected with CLDN4 (C) Western blot showed that the expression of MMP2 and MMP9 was down-regulated in HEp-2 cells transfected with CLDN4 (D) Histogram results of MMP2 and MMP9 protein expression in HEp-2 cells transfected with CLDN4 (E) Western blot showed that the expression of p-JNK increased significantly when the concentration of anisomycin was 2μM. (F) The histogram showed that when the concentration of anisomycin was 2μM, the expression of p-JNK increased most significantly. (G) Western blot showed that the expression of p-JNK increased significantly when anisomycin was treated for 24 hours. (H) The histogram suggests that when anisomycin acts for 24 hours, the expression of p-JNK increases most significantly.

### 2.4 Effect of JNK activator AM (anisomycin) on the expression of p-JNK, JNK, MMP-2, and MMP-9

To confirm the effect of CLDN4 expression on p-JNK, MMP-2, and MMP-9 expression, we treated HEp-2 cells with the JNK activator AM (aniosmycin). To determine the optimal concentration of AM, HEp-2 cells overexpressing CLDN4 were treated with AM at concentrations of 0.5, 1, 2, 4, and 8 μM for 48 hours. Western blotting showed that, at 2 μM AM, the expression of p-JNK increased significantly, but that of JNK did not change. Therefore, 2μM was used for subsequent testing (Figures 4 E and F). Next, to determine the duration of AM treatment, HEp-2 cells overexpressing cldn4 were treated with 2 μM AM for 6, 12, 24, 36, or 48 hours, and Western blotting showed that while the expression of p-JNK increased significantly at 24 h (P < 0.05), that of JNK did not change significantly. Therefore, 24 hours was optimal (Figure 4 G and H). Next, we divided HEp-2 cells into the following four groups, viz., AM-treated no-load experimental group (VEC + AM), AM-treated overexpression CLDN4 experimental group (CLDN4 + AM), no-load control group (VEC), and overexpression CLDN4 control group (CLDN4), and analyzed the expression of p-JNK, JNK, MMP-2, and MMP-9 by Western blotting. We show that the expression levels of p-JNK, MMP-2, and MMP-9 in both groups of AM-treated cells were significantly higher than those seen in the control groups. Specifically, compared to the “VEC” group, the expression of p-JNK, MMP-2, and MMP-9 was significantly upregulated in the “VEC + AM” group. Compared to the “CLDN4” group, the expression of p-JNK, MMP-2, and MMP-9 was significantly upregulated in the “CLDN4 + AM” group. Compared to the “VEC” group, the expression of p-JNK, MMP-2, and MMP-9 was significantly downregulated in the “CLDN4” group and compared to the “VEC + AM” group, the expression of p-JNK, MMP-2, and MMP-9 was significantly downregulated in the “CLDN4 + AM” group (Figure 5 A-D). These results suggest that AM activates JNK, which enhances the expression of its downstream targets, MMP-2 and MMP-9.

**Figure 5.**
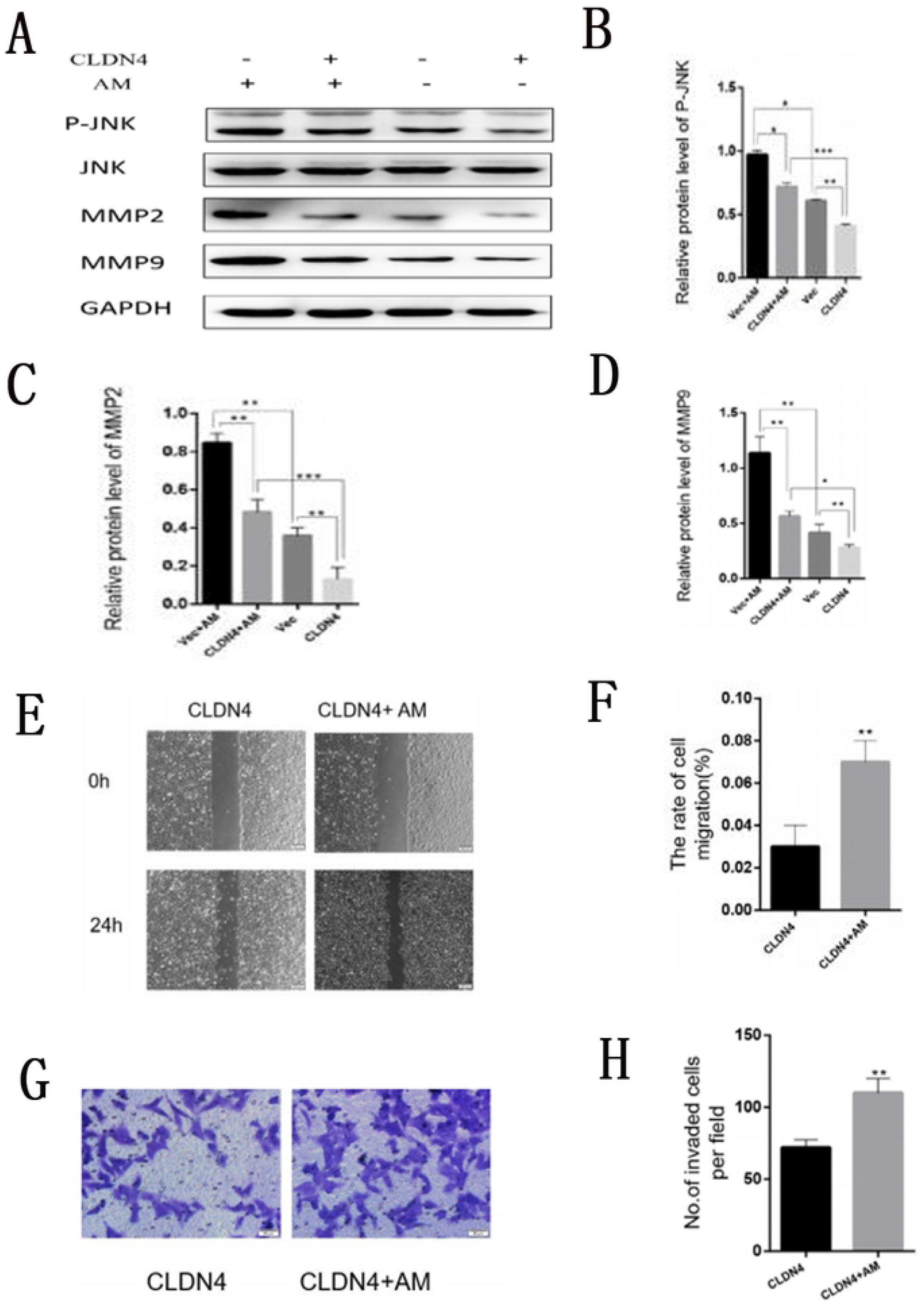
(A) The expression levels of p-JNK, JNK, MMP2 and MMP9 were detected by Western blot (B-D) Columnar statistical chart of the expression levels of p-JNK, MMP2 and MMP9 in four groups of cells(E-F) The migration ability of HEp-2 cells transfected with CLDN4 was enhanced by AM treatment (G-H) The invasive ability of HEp-2 cells transfected with CLDN4 was enhanced by AM treatment.

### 2.5 Effect of the JNK activator AM on migration and invasiveness of laryngeal cancer cells

Next, CLDN4-transfected HEp-2 cells were treated with AM, and its effect on cell migration was evaluated using the scratch test. Compared to untreated cells, AM treatment significantly enhanced cell migration (P < 0.05; Figure 5 E and F). Further, Transwell experiments showed that the invasiveness of cells in the AM-treated group was significantly higher than that of the untreated cells (P < 0.05; Figure 5 G and H).

## Discussion

The human claudin family contains 28 members closely related to transmembrane proteins and is associated with various cancers [15]. Abnormal expression of CLDN4 appears in ovarian cancer [16], esophageal cancer [17], breast cancer [18], and other tumors, but studies on CLDN4 and laryngeal cancer are rare. We previously confirmed that CLDN4 expression is downregulated in laryngeal squamous carcinoma tissue, that it is negatively correlated with methyl-CpG-binding protein 2 expression, and CLDN4 is hypermethylated in HEp-2 cells [4]. As binding of the transcription factor Sp1 is inhibited by mCpmCpG methylation, blocking Sp1 binding will initiate de novo methylation of the CpG islands, leading to promoter silencing [6]. Additionally, in ovarian cancer cells, the CLDN4 promoter is controlled by epigenetic modifications of the Sp1-containing critical promoter region [19]. Similarly, we demonstrate a correlation between the expression of Sp1 and CLDN4 in laryngeal squamous cell carcinoma, along with hypermethylation in the promoter region of CLDN4 gene. Based on these observations, we speculated that there might be a Sp1 binding site in the promoter region of the CLDN4 gene in laryngeal cancer cells and, therefore, treated HEp-2 cells with a Sp1 transcription inhibitor, MTM, which significantly reduced the expression of CLDN4 mRNA and protein.

c-Jun N-terminal kinases (JNKs) belong to the MAPK family. They are involved in various cellular processes, including cell proliferation, differentiation, migration, inflammation, and apoptosis [20,21]. However, the role of JNK in cancer remains controversial, and there is some data to suggest that JNK, especially JNK1, contributes to both malignant transformation and tumor growth [22]. Chang et al. showed that combined treatment of Cal-27 cells with EGCG and gefitinib could phosphorylate JNK, thereby inhibiting tumor cell metastatic potential [23]. In contrast, Chuang et al. have pointed out that IL-6 enhances the migration of oral cancer cells by upregulating the Syk/JNK/AP-1 signaling pathway and that microRNA-196 (miR-196) activates phosphorylated JNK and MMP1/9, thereby enhancing cell migration and invasion [24,25]. Hong et al. have suggested that a change in CLDN4 expression in cancer cells may affect their proliferation by regulating Rho-GTP and p-JNK pathways [17]. Geetha et al. have reported that osmotic stress-induced tight junction disruption was associated with the Tyr-phosphorylation in occludin, ZO-1, and CLDN4 and that inhibition of JNK abrogated such stress-induced Tyr-phosphorylation [26].

The Phospho-SAPK/JNK (p-JNK) antibody does not recognize unphosphorylated SAPK/JNK, although it may slightly cross-react with phospho-Erk1/2 or -p38 that are phosphorylated at homologous residues. It will also react with SAPK/JNK, which is singly phosphorylated at Thr183 or at Tyr185. We show that the expression of CLDN4 and p-JNK is significantly negatively correlated in laryngeal carcinoma cells, that the migratory and invasive abilities of HEp-2 cells transfected with CLDN4 were significantly decreased, and that the expression of p-JNK was significantly downregulated in these cells. Therefore, we speculate that the expression of CLDN4 in laryngeal cancer cells may be related to the JNK signaling pathway.

Matrix metalloproteinases (MMPs) are proteolytic enzymes that degrade the extracellular matrix and thereby facilitate invasion [27]. MMPs can be classified as collagenases, gelatinases, stromelysins, matrilysins, and MT-MMPs. The most common are the collagenases, including MMP-2 and MMP-9. Multiple studies have demonstrated an association between MMP-2 activity levels, cellular invasion, and disease severity in gliomas [28,29,30]. Angela et al. have suggested that wnk2 promoter methylation and silencing in gliomas are related to JNK activation, MMP-2 expression, which may lead to changes in the invasive ability of tumor cells [27]. Studies have also shown that the translationally controlled tumor protein can increase the secretion of MMP-9 and the translocation of p-JNK from the cytoplasm to the nucleus. Further, silencing of JNK and MMP-9 significantly inhibited the invasion of rhTCTP-enhanced CRC cells into Matrigel [31]. These observations suggest that MMPs may be associated with the JNK signaling pathway and lead to changes in tumor invasiveness.

We report a positive correlation among p-JNK, MMP-2, and MMP-9 in laryngeal squamous cell carcinoma. Additionally, transfection of CLDN4 resulted in the downregulation of p-JNK in HEp-2 cells, along with a significant decrease in the expression of MMP-2 and MMP-9 proteins, suggesting an association between the downregulation of p-JNK and the expression of MMP-2 and MMP-9. AM is a JNK and p38 activator that inhibits the formation of linear staining of ZO-1. Thus, inhibition of JNK restored linear staining of ZO-1 after AM treatment, but CLDN 4, 6, and 7 could not fully colocalize at the ZO-1 positive sites [32]. To confirm this relationship between the JNK signaling pathway and MMP-2 and -9, we treated HEp-2 cells with AM. We found that p-JNK levels increased significantly, which then enhanced the expression of MMP-2 and -9. Fei et al. reported that an increase in CLDN4 expression due to PAK4-mediated CEBPB activation could promote breast cancer cell migration and invasion [33], and CLDN4 can promote the invasion of ovarian cancer cells by activating MMP-2 [34]. It can also enhance the proliferation and invasion of gastric cancer cells [35]. Moreover, CLDN4-mediated destruction of the TJ enhances the invasiveness and metastasis of cancer cells in human CRC [36]. While these studies establish an important role for cldn4 in the migration and invasion of cancer cells, the specific mechanism remains unclear. We also confirm that changes in cldn4 expression can affect the migration and invasion of laryngeal squamous cell carcinoma, report that this may be related to the activation of JNK signaling pathway, and show that MMP-2 and MMP-9 are involved in this process. To summarize, our results show that the promoter region of CLDN4 gene is hypermethylated in laryngeal squamous carcinoma cells. That inhibition of the transcription factor Sp1 can lead to the downregulation of CLDN4 expression. Transfection of CLDN4 into HEp-2 cells inhibited their migration and reduced their invasiveness, significantly downregulated the expression of p-JNK, and decreased the expression of MMP-2 and MMP-9 proteins, suggesting that this process may be related to the JNK signaling pathway.

## Methods

### Materials and Methods

#### Patient samples and cell culture

This study used tissue samples from 84 cases of laryngeal squamous cell carcinoma and 76 cases of adjacent tissues obtained from the Department of pathology, First hospital of Jilin University.

#### Immunohistochemistry

Immunohistochemical staining was performed to analyze the expression of CLDN4, SP1, P-JNK, MMP-2, and MMP-9 in LSCC and adjacent tissues. Tissue sections were incubated at 70°C for 1 h, followed by incubation in 100% xylene for 10 min, 100% ethanol twice for 5 min each, and then 95%, 90%, and 80% ethanol for 2 min each. Slides were washed and fixed under high pressure. An automatic immunohistochemical staining program was used to counterstain, differentiate, anti-blue stain, dehydrate, and clear the slides, which were then observed under a microscope.

#### Cell culture and cell transfection

HEp-2 cells and HeLa cells were provided by the Key Laboratory of Pathology and Biology, Ministry of Education, Jilin University. The cells were cultured in H-DMEM containing 10% fetal bovine serum in an incubator with 5% CO_2_ at 37°C. 5-aza-dC was prepared in DMSO; specifically, 5-aza-dC powder was dissolved at a concentration of 200 M and stored at −20 °C. Control and experimental groups were established as follows. After conventional cell culture for 24 hours, HEp-2 cells in the experimental group were treated with 5-aza-dC for 24 hours. According to the manufacturer’s instructions, cell transfection was performed using the Lipofectamine 3000 kit (Invitrogen, Carlsbad, CA, USA), and the cells were harvested 48 hours after transfection.

#### Bisulfite genomic sequence (BSP)

DNA from cells in the experimental and administration groups was sequenced as follows. Genomic DNA was extracted from HEp-2 cells, and primers were designed for the promoter region of the cldn4 gene, which PCR, modified by sodium bisulfite, then amplified, and 10 positive clones were selected for sequencing. Sequencing results were analyzed, the methylation ratio of the CpG sites was calculated, and the distribution map of the methylation beads was drawn.

#### RT-PCR

Total RNA was extracted using the standard phenol-chloroform extraction and isopropanol precipitation method. All cDNAs were prepared according to the manufacturer’s instructions with 1 μg of RNA and the SuperScript II enzyme (Monad, Shanghai, CHINA). One-tenth to one-twentieth of this reaction volume was used as a template for PCR amplification with Taq polymerase (Monad). Amplification was carried out for 30 cycles at 95 °C for 1 min, 55 °C for 1 min, and 72 °C for 2 min. Primer sequences were identical to those reported in ref. 4.

#### Western blotting

Hep-2 cells were washed with cold phosphate-buffered saline (PBS), and the culture dishes were placed on ice to extract protein. Cells were lysed with 500 μL of lysate per dish for 30 min, cells scraped into Eppendorf tubes, and centrifuged at 150 rpm for 15 min. The supernatant was retained and used for measuring protein concentration. For electrophoresis, 30 μg of the protein sample was added to each well and electrophoresed at 80 V for 30 min and at 120 V for 90 min. Proteins were then transferred to a membrane and incubated sequentially with primary and secondary antibodies.

#### Transwell invasion assay

According to the manufacturer’s instructions, the Transwell invasion assay was performed using Transwell plates (Corning, New York, USA). Cells (∼ 5 × 10^4^) were added to the upper compartment of the culture chamber. After 24 hours of incubation at 37°C with 5% CO_2_, the number of cells invading through the Matrigel matrix was calculated in 10 visual fields that were randomly selected from the central and peripheral portions of the filter using an OLYMPUS microscope.

#### Scratch test

The cells were inoculated into 6-well plates, and on the second day, a sterile pipette gun head was used to a scratch perpendicular line in the 6-well plate to produce a straight wound. The wound area was cleaned thrice with PBS and was imaged under a microscope at 0 and 24 hours after wounding. The parallel distance between the scratch lines was observed, measured under the microscope at 0 h, and recorded as a value “a”. After 24 h, images were acquired at the same position, and the parallel distance was measured and recorded as value “b”. Mobility was calculated using the formula (a − b) / a × 100 (%).

### Statistical analysis

Spearman’s correlation was used to analyze the correlation between CLDN4 and SP1 expression among CLDN4, p-JNK, MMP-2, MMP-9, and between p-JNK and MMP-2 and MMP-9.

## Abbreviations

CLDN4: Claudin 4
MTM: Mithramycin A
JNK: c-Jun N-terminal kinase
SP1: specifieity protein 1
MMP: matrix metalloproteinases
AM: anisomycin

## Acknowledgments

None

## Author Contributions

Jixuan Liu and Yafang Liu were involved in drafting and final revision of the manuscript. Yafang Liu, Kaiyue Peng and Ya Liu were involved in designing and completing the experiment.

## Ethics declarations

The patient provided informed consent, and the article was approved by the Ethical Committee of the First Hospital of Jilin University in Changchun, China. We obey the principles of the 1983 Declaration of Helsinki. In other words, all of experiments in this paper obey this principle.

